# VARUS: Sampling Complementary RNA Reads from the Sequence Read Archive

**DOI:** 10.1101/608737

**Authors:** Mario Stanke, Willy Bruhn, Felix Becker, Katharina Hoff

## Abstract

Vast amounts of next generation sequencing RNA data has been deposited in archives, accompanying very diverse original studies. The data is readily available also for other purposes such as genome annotation or transcriptome assembly. However, selecting a subset of available experiments, sequencing runs and reads for this purpose is a nontrivial task and complicated by the inhomogeneity of the data.

This article presents the software VARUS that selects, downloads and aligns reads from NCBI’s Sequence Read Archive, given only the species’ binomial name and genome. VARUS automatically chooses runs from among all archived runs to randomly select subsets of reads. The objective of its online algorithm is to cover a large number of transcripts adequately when network bandwidth and computing resources are limited. For most tested species VARUS achieved both a higher sensitivity and specificity with a lower number of downloaded reads than when runs were manually selected. At the example of twelve eukaryotic genomes, we show that RNA-Seq that was sampled with VARUS is well-suited for fully-automatic genome annotation with BRAKER.

With VARUS, genome annotation can be automatized to the extent that not even the selection and quality control of RNA-Seq has to be done manually. This introduces the possibility to have fully automatized genome annotation loops over potentially many species without incurring a loss of accuracy over a manually supervised annotation process.

## Background

A very large amount of next generation sequencing (NGS) data is being deposited in public databases such as the sequence read archive (SRA) of the National Center for Biotechnology Information (NCBI) [1] and the European nucleotide archive (ENA) [2]. At the time of writing, in March 2019, the SRA stored about 2.7·10^16^ bp of raw sequencing data [3]. This archived data provides experimental support for a very large number of manifold individual studies with very specific purposes for diverse species.

A large fraction of this NGS data is RNA-Seq. In many studies, RNA has been sequenced from many replicates to draw statistical conclusions about the expression of transcripts [4, 5]. Besides being a requirement for the reproducability of the original studies, a large value of the repository lies in the possibility to use the data for further studies including such with very different purposes than the original studies. One important reason to repurpose or use RNA-Seq is structural genome (re)annotation. Often a genome is annotated only after several transcriptomic studies of the same species have been conducted. There are many reasons for structural genome *re*annotation as well, e.g. the assembly has been improved, the genomes of further strains have been sequenced, to apply newer more accurate annotation methods, or annotate a large number of species’ genomes consistently.

Taking *all* available RNA-Seq data for a species is often not feasible or counterproductive due to issues of the quality of data or alignments. The available data can be very abundant and redundant. For example, there are more than 30,000 RNA-Seq sequencing runs for *Drosophila melanogaster* (Table 1) totaling to more than 41 tera base pairs. Further, quality control of sequencing runs is necessary as some genomic libraries are mislabeled as transcriptomic, contain adapter sequences or simply have low quality or alignability (see Table 1). Therefore, it is sensible to choose a *subset* of all available reads for genome annotation. In doing this, the *complementarity* of chosen reads should be the aim in order to cover a large fraction of all transcripts of the organism, while keeping the burden on computation and data throughput manageable. Due to tissue- or condition-specific expression it may be necessary to use reads from many different sequencing runs from a diverse set of tissues or conditions. The meta data that the uploaders entered includes often an abstract of the study. The ‘manual’ scan of these abstracts is tedious, is not always conclusive on tissue and condition and – even if so – does not allow to conclude which fraction of the transcripts may be expressed that have not yet been covered by other choices of sequencing runs. Further, the meta data could be incorrect.

**Table 1:** Transcriptomic sequencing runs. The second column shows the total number of RNA runs in SRA for the studied species. The last column shows the percentage of runs that were sampled by VARUS of which the first batch exhibited very low unique alignability, more specifically, at least 95% of reads aligned either not at all or multiple times using HISAT2. Such runs are subsequently ignored by VARUS.

| species | transcriptomic runs in SRA | $\leq 5\%$ of reads align uniquely |
| --- | --- | --- |
| *Anopheles gambiae* | 597 | 15.5% |
| *Bombus terrestris* | 357 | 6.4% |
| *Chlamydomonas reinhardtii* | 1250 | 7.1% |
| *Cucumis sativus* | 685 | 21.2% |
| *Drosophila melanogaster* | 30207 | 22.8% |
| *Fragaria vesca* | 170 | 10.6% |
| *Hymenolepis microstoma* | 31 | 32.2% |
| *Medicago truncatula* | 1252 | 9.2% |
| *Parasteatoda tepidariorum* | 80 | 16.3% |
| *Prunus persica* | 268 | 13.2% |
| *Saccharomyces cerevisiae* | 16139 | 30.5% |
| *Verticillium dahliae* | 30 | 40.0% |

Ohta, Nakazato and Bono performed a large-scale assessment of reported read qualities in SRA, but did not align the reads at all [6]. To our knowledge no previous software existed for automatically sampling reads from SRA from a maximal set of transcripts. A trivial random sampling from all libraries is not optimal as there may be large biases on the tissues or conditions that are represented in the archive.

We present the new tool VARUS [7] that automatically samples, downloads and aligns reads from all runs available on SRA for a given species. It implements an online algorithm that iteratively downloads small random samples from sequencing runs and thereby selects such runs for download of further samples that were estimated to complement best the previously downloaded reads and that have a relatively high percentage of reads or reads pairs that align uniquely to the genome. We tested VARUS on twelve genomes and compared the introns suggested by spliced alignments to those of the reference annotations. Compared with a manual selection of sequencing runs, VARUS achieves for most species a higher accuracy with a smaller number of downloaded reads than the manual selection.

## Implementation

VARUS takes as input binomial (Latin) species names and corresponding genome FASTA files (see Figure 1). A script (`RunListRetriever.pl`) queries the SRA for a list of IDs of all sequencing runs of molecule type *RNA* for the given species. The output, a ‘run list’, is then used by VARUS to choose, download and align samples of RNA-Seq reads. The alignment is performed automatically, either with HISAT2 [8] or STAR [9]. For each species a single BAM formatted file with spliced alignments is output.

**Figure 1:**
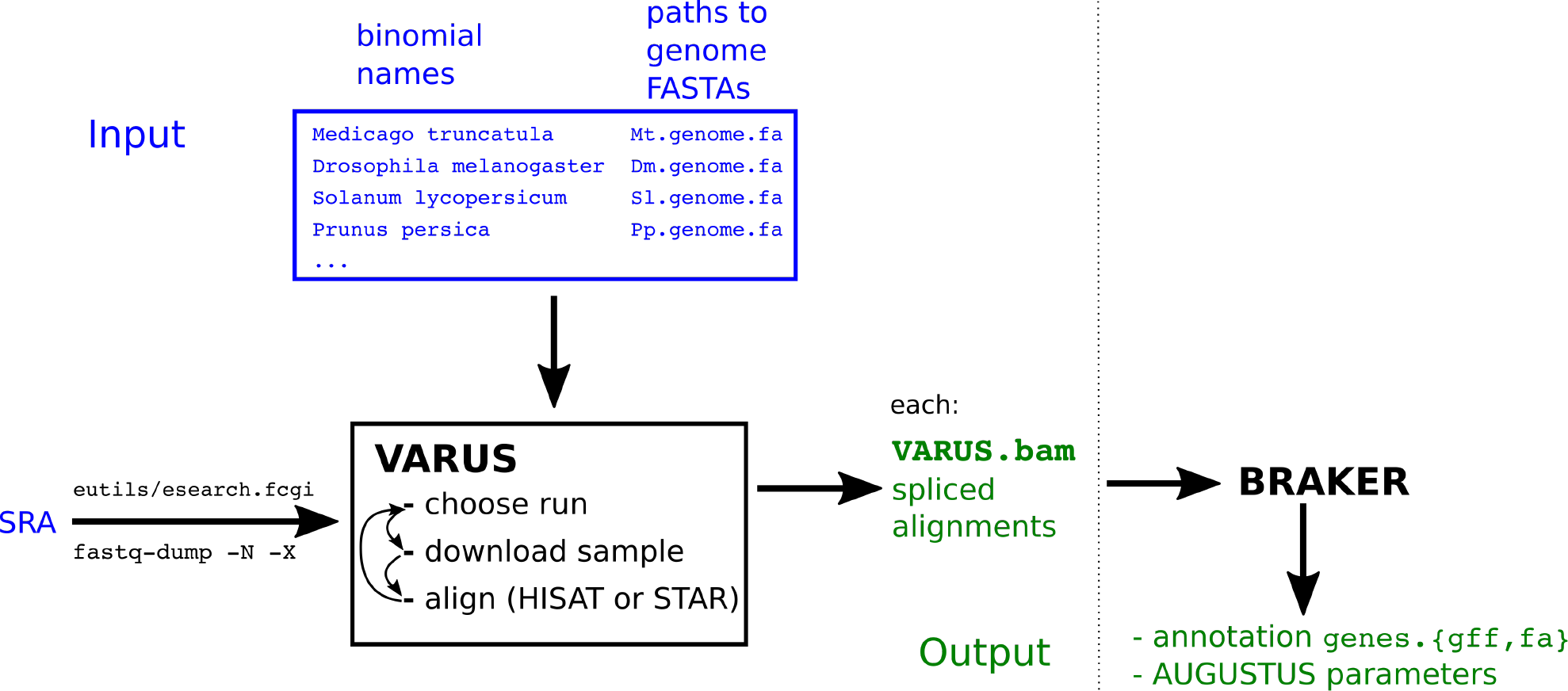
VARUS flowchart. VARUS itself outputs a file `VARUS.bam` with all spliced alignments for each species in the input list. In this study, these alignments were used to annotate the genomes with BRAKER [10].

The objective of VARUS is to collect samples of RNA-Seq reads from all available sequencing runs that together adequately cover as many transcribed RNAs as possible. We neither make prior assumptions on the expression profiles of runs nor do we require an (assembled) transcriptome as input. *Adequate coverage* is a vague formulation and certainly depends on the downstream application. We use the VARUS output for genome annotation, where we expect that the accuracy, with which an individual gene structure can be annotated, increases with its coverage when it is lowly covered, but the expected accuracy returns are diminishing with increasing coverage. Formally, our objective is to maximize a score

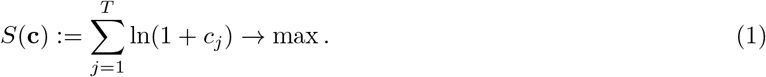

Here, the *transcribed units* are numbered from 1 to *T*, *c*_*j*_ is the count of aligned reads or read pairs from transcribed unit *j* and **c** = (*c*_*1*_, …, *c*_*T*_). The logarithm was chosen to reflect diminishing returns of additional coverage for genome annotation accuracy. As we do not know the set of all transcribed RNAs themselves in our application setting, we use genome tiles (of default size 5Kb) as proxy for transcribed units. Admittedly, such tiles may not be expressed at all or may contain multiple genes. However, the idea is that the similarity of expression profiles from different runs is approximated sufficiently using whole-genome tile coverages for purposes of annotation.

VARUS implements a heuristic online algorithm to approximate above goal step by step. If **c** is the current vector of read counts for all transcribed units, the next batch of *b* reads (default *b* = 50, 000) to download and align are chosen from run

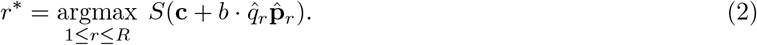

Here, the available runs are numbered from 1 to *R*, 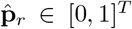 is the parameter vector of a multinomial distribution that is an estimate of the distribution on 1..*T* of a randomly drawn read from the *r*-th run. 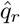 is an estimate of alignment quality, a weighted sum of the fraction of uniquely alignable and spliced reads. Ties occur – in particular for all runs of which no reads have been downloaded yet at all – and are broken randomly with a bias towards longer average read lengths. In (2) the next batch is chosen greedily from a run so as to maximize the score of the expected count vector after the choice, 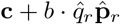.

When a run for downloading the next batch of reads has been chosen, VARUS calls `fastq-dump` which allows to download a range of consecutive ‘spots’ from a specified run. VARUS keeps track of the set of all batches of a run that have been downloaded previously, so that eventually all reads would be downloaded exactly once. However, VARUS downloads (at most) only a fixed number of batches (here mostly `maxBatches`=1000).

## Results

### Data Sets

We tested VARUS on the twelve species listed in Table 1, which include model organisms with much data as well as other organisms with relatively little data. Table 1 also shows that 6%-40% of sequencing runs do not satisfy our requirements for quality: At least 5% of reads needed to align uniquely to the genome using HISAT2. Paired libraries were thereby handled accordingly. For command lines see Supplementary Material. The reasons for failing even these weak criteria are diverse. They include the insufficiency of the alignment program for the sequencing platform, but also inappropriate data for that purpose such as miRNA sequencing runs, a mislabeled source species and very short reads. An example of the latter is *S. cerevisiae*, where 13% of the sampled runs had at least one out of two (‘paired’) sequence files with an average read length of ≤ 10 base pairs. Further we found cases where genomic libraries were clearly mislabeled as transcriptomic.

We compared VARUS to a selection procedure in which a person browses the SRA archive and selects and chooses sequencing runs for download. We refer to this as the *manual* selection method. Afterwards the alignment against the genome is performed in the same way as in VARUS. Sequencing runs that appeared to be of low alignability were discarded and not counted towards the number of downloaded reads. This is to the benefit of the manual method as all downloaded reads are counted in the VARUS analysis. The manual selection gave preference to long paired reads from the Illumina platform (e.g. 2 × 150 bp) that were from different experiments and different tissues. The interpretation of the natural language description of the experiment in the meta data can be difficult and a manual classification to tissue or condition would not be highly repeatable. In Figure 2 we show as an example, a visualization of expression diversity with a tissue subset (*Drosophila* gut) highlighted. A systematic approach to select maximally complementary sets of tissue types or conditions from meta data alone does not seem possible. Rather the data itself needs to be examined.

**Figure 2:**
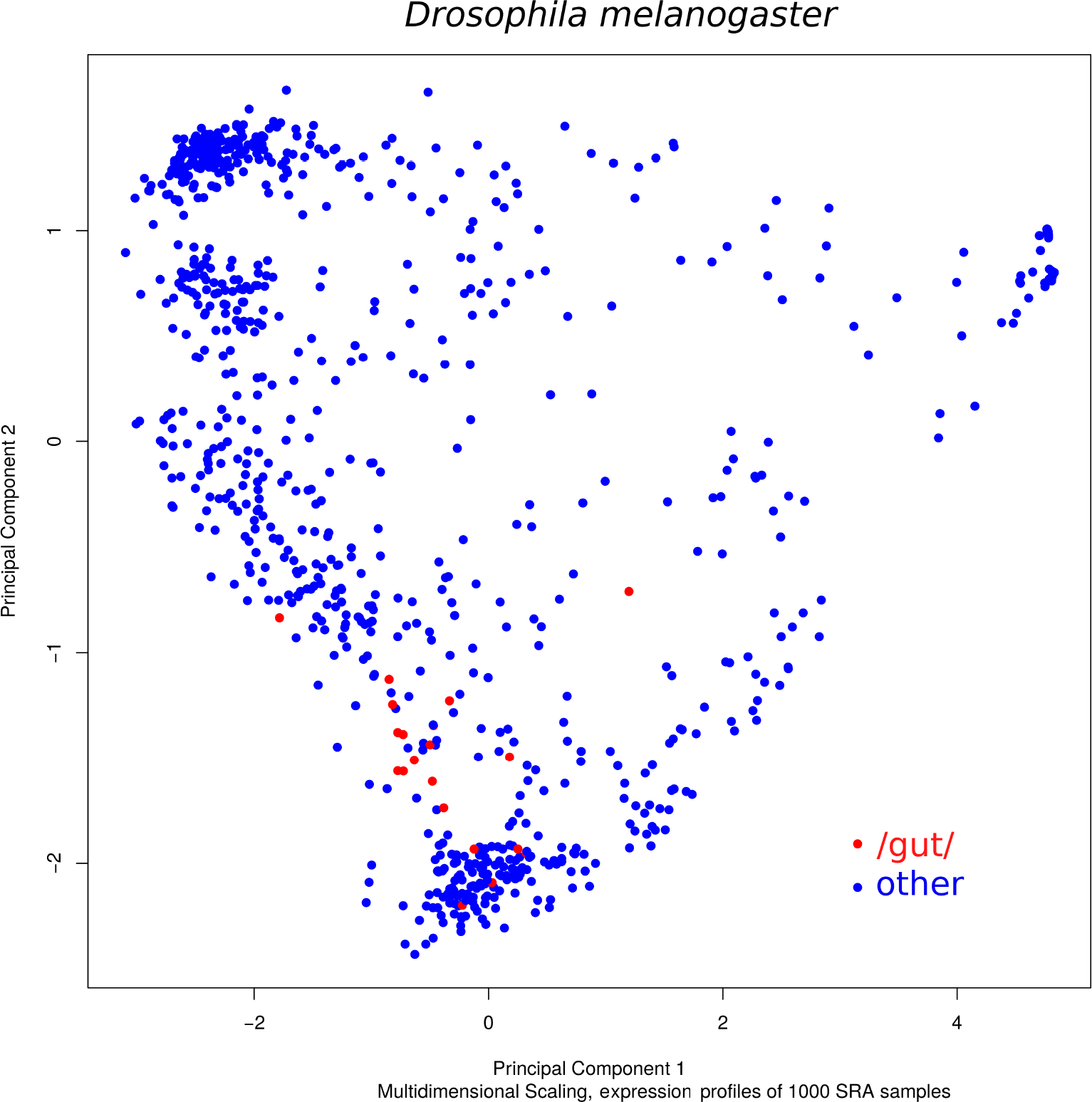
Expression diversity. Multidimensional scaling plot of the expression profiles of 1000 runs sampled from *Drosophila melanogaster* from SRA. The red dots mark those runs, whose meta data description included the search string “gut”. The plot was created with edgeR [5].

### Accuracy

To evaluate and compare the performance of the read selection methods, we measured the accuracy with which the *introns* of a reference annotation (see Supplementary Table 2, reference annotations were pre-processed with GenomeTools [11]) are found with spliced alignments of the selected reads. This intron accuracy measure is a proxy to the relative performances that could be achieved when using the alignments for genome annotation. The intron accuracy measure does not depend on any genome annotation method that ought to be evaluated and is therefore a more direct performance measure. As a reference, we take for each genome the set of introns in the protein-coding regions of genes. The annotation of such ‘coding’ introns is usually more reliable that of introns in untranslated regions or non-coding genes. As a prediction, we take the set of introns *induced by the spliced alignments* of the selected reads. In this context an intron is a pair of genome coordinates and each intron is counted only once, i.e. the sets are defined in the usual mathematical sense. The *sensitivity* and *specificity* are defined as usual in the genome annotation context: The (intron) *sensitivity* is the percentage of reference introns that were also predicted. The (intron) *specificity* is the percentage of predicted introns that were also in the reference set.

**Table 2:** Intron accuracy. The sensitivity (sn) and specificity (sp) with which VARUS and the manual method find introns in the reference genome annotation. The last two columns shows the number of reads or read pairs (spots) in millions that have been downloaded from SRA and from how many different runs they stem. Better values are typeset in boldface.

|  |  | intron |  | #spots | # seq. |
| --- | --- | --- | --- | --- | --- |
|  |  | sn | sp | [M] | runs |
| *Anopheles gambiae* | VARUS | **.805** | **0.23** | 50 | 566 |
|  | manual | .269 | .179 | **18.4** | 5 |
| *Bombus terrestris* | VARUS | **.918** | .365 | 50 | 357 |
|  | manual | .666 | **.623** | **29.1** | 3 |
| *Chlamydomonas reinhardtii* | VARUS | **.687** | **.37** | **50** | 618 |
|  | manual | .68 | .26 | 88 | 5 |
| *Cucumis sativus* | VARUS | **.96** | **.414** | **50** | 643 |
|  | manual | .91 | .262 | 126.6 | 7 |
| *Drosophila melanogaster* | VARUS | **.935** | **.359** | **50** | 758 |
|  | manual | .896 | .264 | 58.5 | 5 |
| *Fragaria vesca* | VARUS | **.915** | **.475** | **50** | 170 |
|  | manual | .823 | .322 | 71.9 | 7 |
| *Hymenolepis microstoma* | VARUS | **.886** | .237 | 50 | 31 |
|  | manual | .778 | **.391** | **15.7** | 2 |
| *Medicago truncatula* | VARUS | .717 | **.386** | **50** | 403 |
|  | manual | **.723** | .21 | 214.9 | 6 |
| *Parasteatoda tepidariorum* | VARUS | **.869** | **.469** | **50** | 80 |
|  | manual | .83 | .352 | 71.9 | 6 |
| *Prunus persica* | VARUS | **.945** | **.188** | 400 | 266 |
|  | manual | .942 | .138 | **328.8** | 6 |
| *Saccharomyces cerevisiae* | VARUS | **.864** | **.004** | **50** | 983 |
|  | manual | .85 | .002 | 99.8 | 5 |
| *Verticillium dahliae* | VARUS | **.681** | .222 | **50** | 30 |
|  | manual | .662 | **.336** | 81.9 | 3 |

Figure 3 shows for the examples of *Drosophila melanogaster* and *Fragaria vesca* how the sensitivity grows and the specificity changes as the number of reads or read pairs (*spots* in the language of SRA) grows. A relatively ‘small’ number of spots, 3 million, is in all twelve species sufficient to cover with VARUS at least half of the introns (data not shown). For both species, VARUS achieves the same sensitivity as the manual selection with fewer spots, in the case of strawberry even with a small fraction of the spot number. Further, when the sensitivity of VARUS and manual selection are equal, the specificity of VARUS is much higher than the specificity of the manual selection. As can be expected, the intron sensitivity initially grows fast and eventually saturates. When the number of aligned reads grows, false positive introns are accumulated at an increasingly higher rate than true positives. They can include alignment errors and very minor nonfunctional splice forms and could deteriorate annotation accuracy if left unfiltered.

**Figure 3:**
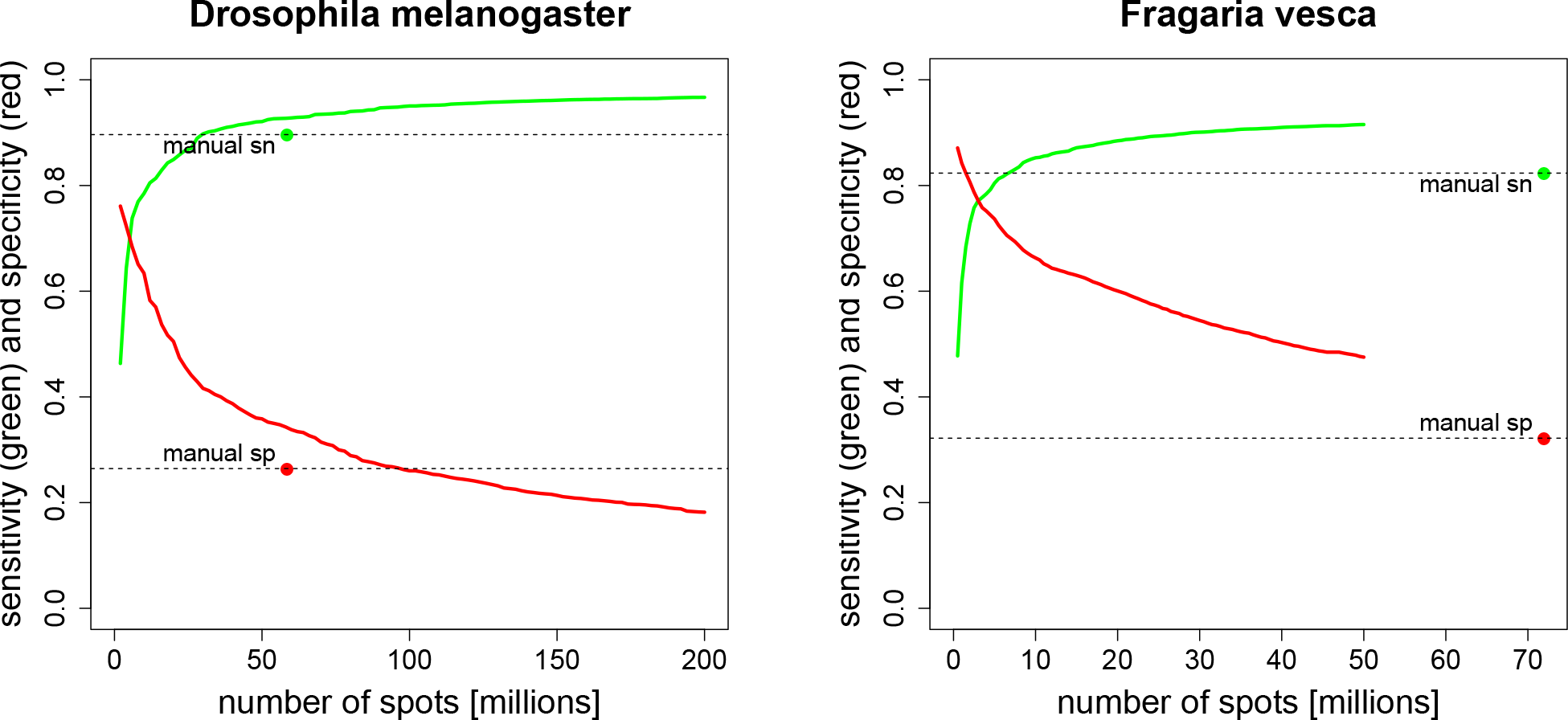
Intron Accuracy. VARUS’ sensitivity (green curve) and specificity (red curve) as a function of the downloaded number of spots (= reads or read pairs), in order of download. The manual selection method (green and red dot) has a fixed number of reads and horizontal dashed lines at manual sensitivity and specificity are drawn to facilitate comparisons.

Table 2 shows the accuracies and download numbers for all examined species. The sensitivity and specificity curves for the remaining species (like in Figure 3) are included in the Supplementary Materials. For all species but *Verticillium dahliae* VARUS achieved both a higher sensitivity and specificity with a lower number of downloaded reads or read pairs than the manual selection of whole runs. For *Chlamydomonas reinhardtii*, *Cucumis sativus*, *Drosophila melanogaster*, *Fragaria vesca*, *Parasteatoda tepidariorum* and *Saccharomyces cerevisiae* this can be seen directly from Table 2. For *Anopheles gambiae*, *Bombus terrestris*, *Hymenolepis microstoma*, *Prunus persica* VARUS downloaded more reads or read pairs than the manual methods, but the curves of sensitivity and specificity as a function of cumulatively downloaded reads (Supplementary Figure 1) show that VARUS had a higher specificity and a lower number of downloaded reads or read pairs when the sensitivity first surpassed that of the manual method (horizontal dashed line through ‘manual sn’). For *Verticillium dahliae* the two methods are not objectively comparable unless one weights computational resources against accuracy. When the sensitivities of the two methods were equal (66.2%), VARUS’ specificity was slightly lower than that of the manual selection (31.5% versus 33.6%), but it had downloaded less than one third of the number of spots (25.5 million versus 81.9 million).

### Whole-Genome Annotation

We used the selected RNA reads for genome annotation and examined their relative influence on accuracy. For this we used the fully automatic genome annotation pipeline BRAKER [10]. BRAKER obtained as input only the intron candidates derived from the spliced alignments of the reads against the genome as ‘hints’ and the repeat-masked genome. No other input or previous parameters are used by BRAKER. The BRAKER pipeline uses the hints to compile a training set of gene structures with GeneMark-ET [12] and then AUGUSTUS [13, 14] to structurally annotate the whole genome using again the intron hints.

Figure 4 shows a comparison between the gene prediction accuracies of BRAKER (obtained with Eval [15]) when either the manually selected runs were used or the reads selected by VARUS. On average, VARUS downloaded fewer reads (Figure 4, left) than were downloaded manually. Yet, with VARUS the whole-genome annotation accuracy tends to be rather higher than with the manual selection. As can be seen at the right side of Figure 4, in 8 of the species, the results with VARUS are better than with the manual selection, in the remaining 4 species the manual selection is better. However, the clear advantage of VARUS in terms of intron accuracy does not carry over to a clear annotation accuracy advantage with BRAKER. This suggests that BRAKER may not be fully exploiting the additional accuracy or that most of the additional true positive introns would have been found anyway and that most of the false positive introns would have been filtered out anyway. We conclude that VARUS automatizes a task that hitherto needed manual work at even a small average benefit to the accuracy of genome annotation.

**Figure 4:**
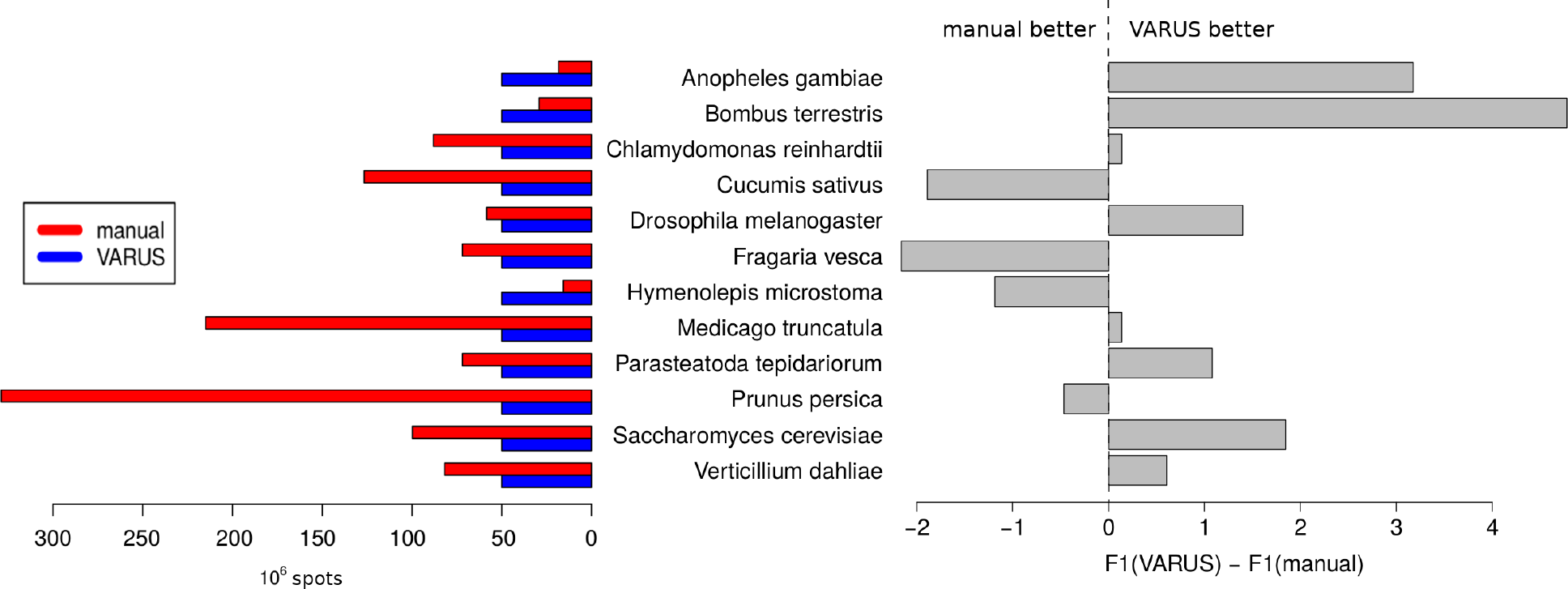
Relative Annotation Accuracy. The right side shows the difference in whole-genome annotation accuracy in percent. The F1-measure of coding exon accuracy (the harmonic mean of sensitivity and specificity) was used, whereby either annotation was compared to the respective reference annotation (see Supplementary Table 2). The left shows the corresponding input data set sizes. The average and mean number of spots chosen by the manual method are 100 and 77 million, respectively, and larger than the 50 million spots downloaded by VARUS. Averaging over the species, the F1 accuracy of BRAKER is 0.62% higher when RNA-Seq is selected by VARUS rather than manual.

## Discussion

VARUS provides a convenient way to sample RNA-Seq data from a given species. Even a trivial independent sampling of a random subset of many or all sequencing runs can be very useful, e.g. to estimate and compare library qualities on the basis of the actual data. Such a trivial sampling can be done with VARUS with the option `--loadAllOnce` with which a single batch from each run will be downloaded once. VARUS – even if used in such a trivial way – can decrease the risk that a manual sample of a small number of (complete) sequencing runs bears. It could be an *unlucky* choice of libraries or conditions or tissues that leaves a relatively large fraction of transcripts or major splice forms insufficiently represented.

When more data is available than can be handled reasonably, VARUS allows to achieve a near optimal sensitivity with a small fraction of data. For example, the alignments found by VARUS covered 96.7% of all annotated *Drosophila melanogaster* introns with less than 1/2000 of the total spots available (200 million out of 433 billion spots).

VARUS makes it possible to annotate a large number of species on the basis of RNA-Seq in a loop. To demonstrate this, we used the BRAKER pipeline, that can be trained and run automatically with RNA-Seq and a genome only. The annotation loop does not require manual work, except for compiling a list of species names and genome files as shown in Figure 1. Nevertheless, the overall performance of VARUS+BRAKER appears to be on par or rather better than when reads are selected manually.

We think that VARUS could be useful for transcriptome assembly as well, whether genome-guided transcriptome reconstruction or even *de novo* transcriptome assembly methods. Like genome annotation, these tasks also seek to construct a set of transcripts that is as complete as possible. On the other hand, subsetting the data may be a necessity and the complementarity of the subset is desirable. In addition, Pertea et al. identify large variations in expression levels as a problem for transcriptome assembly [16], while the objective (1) of VARUS gives a relative disincentive to collect more data from transcript units that are already highly represented. For this reason we plan to allow VARUS to make unspliced alignments as well as a future development.

## Conclusions

We introduced the software VARUS to efficiently and automatically collect samples from SRA to cover a large set of transcripts. VARUS makes it easy to repurpose SRA resources for genome annotation without requiring that meta data on the experiments is manually read or automatically mined. We expect that a lot can be gained by even more automatization towards a more continuous process of genome annotation. Annotations could be updated when a new assembly or significantly more RNA-Seq becomes available or when annotation methods are substantially improved. In this context, it should be stressed that while most genome annotation pipelines heavily rely on RNA-Seq data, they often also use other data, such as protein sequences from related species. It can therefore be advisable to update an annotation of one species because of a change in the available data or gene structures of another species.

At the moment, the methods with which the current assemblies of eukaryotic genomes were annotated are quite diverse, e.g. in terms of programs but also in usages and versions of the same program. Such an inhomogeneity can be very relevant for questions of comparative genomics in which the (often rare) *differences* between the genomes are in the focus. Then spurious differences from inhomogeneous annotation methods can dominate the reported differences. For such comparative purposes, the consistent reannotation of all genomes is desirable but may only be practically feasible when no manual work is required for each species.

## Availability and requirements

- Project name: VARUS
- Project home page: https://github.com/Gaius-Augustus/VARUS
- Operating system(s): Linux
- Programming language: C++, Perl
- Other requirements: samtools [17], fastq-dump, HISAT2 or STAR, filterIntronsFindStrand.pl (BRAKER)
- License: GPLv3

## Author’s contributions

MS conceived the problem and algorithm, implemented VARUS, performed VARUS experiments and wrote the manuscript. WB implemented VARUS and performed VARUS experiments. FB performed the manual selection and download of sequencing runs. KJH performed alignments of manually selected libraries, ran BRAKER and wrote the manuscript. All authors read and approved the final manuscript.

## Acknowledgements

The research was supported by the Deutsche Forschungsgemeinschaft (DFG) grant number STA 1009/12-1 to MS and by the US National Institutes of Health grant GM128145 to MS.

