## Supplementary Material for "VARUS: Sampling Complementary RNA Reads from the Sequence Read Archive"

**2 Supplementary Figures** **3**

**3 Supplementary Tables** **5**

### 1 Supplementary Methods

#### 1.1 Assembly & Reference Annotation Processing

The names of genome FASTA file entries downloaded from NCBI are long and complex. For our purposes, unique sequence IDs in the header are sufficient. We trimmed FASTA headers and replaced dots in sequence names by underscores:

```
cat original.fa | perl -pe 's/(>\S*)\.(\d+)\s.*/$1_$2/g;' > genome.fa
```

Dots in sequence names were replaced by underscores in the corresponding annotation files:

```
cat original.gff | perl -pe 's/\./_/' > annot.gff3
```

BRAKER is by design unable to predict genes with frameshift errors or genes that have parts located on both strands. Reference annotation files were checked for genes with such issues using GenomeTools [Gremme et al., 2013]:

```
gt  gff3  -force  -tidy  -o annot_tidy_by_GenomeTools.gff3  \
  -retainids  -sort  annot.gff3  2>  errors_by_GenomeTools
cat errors_by_GenomeTools | grep -v gbunit | cut -f6 -d' ' | sort | uniq -c | wc -l
```

Reference annotation files in GFF3 format were converted to GTF format using GenomeTools (this also removed numerous file entries that are not related to the structures of protein coding genes, e.g. rRNA features, etc.):

```
gt  gff3_to_gtf  -force  -o annot_by_GenomeTools.gtf annot_tidy_by_GenomeTools.gff3
```

Finally, GTF format files were run through validate_gtf.pl from the Eval package [Keibler and Brent, 2003]:

```
validate_gtf.pl -f annot_by_GenomeTools.gtf
```

The resulting file annot_by_GenomeTools.fixed.gtf was used for measuring gene prediction accuracy of BRAKER.

#### 1.2 Running VARUS

VARUS was run with the command

```
runVARUS.pl --readFromTable species.txt --aligner=HISAT > runVARUS.log 2> runVARUS.err
```

Here, species.txt is a tab or semicolon separated file with the binomial names and paths to the genome files:

```
Anopheles gambiae           Anopheles_gambiae.fa
Bombus terrestris           Bombus_terrestris.fa
Chlamydomonas reinhardtii   Chlamydomonas_reinhardtii.fa
Cucumis sativus             Cucumis_sativus.fa
Medicago truncatula         Medicago_truncatula.fa
Drosophila melanogaster     Drosophila_melanogaster.fa
Fragaria vesca              Fragaria_vesca.fa
Parasteatoda tepidariorum   Parasteatoda_tepidariorum.fa
Prunus persica              Prunus_persica.fa
Hymenolepis microstoma      Hymenolepis_microstoma.fa
Saccharomyces cerevisiae    Saccharomyces_cerevisiae.fa
Verticillium dahliae        Verticillium_dahliae.fa
```

In the same directory was a text file VARUSparameters.txt with the following content. runVARUS.pl reads its parameters from there:

```
--batchSize 50000
--maxBatches 1000
--blockSize 5000
--cost 0.001
--deleteLater 0
--estimator 2
--exportObservationsToFile 1
--exportParametersToFile 1
--fastqDumpCall fastq-dump
--genomeDir ./genome/
--lambda 10.0
--loadAllOnce 0
--mergeThreshold 10
--outFileNamePrefix ./
--pathToParameters ./VARUSparameters.txt
--pathToRuns ./
--pathToVARUS /home/mario/VARUS/Implementation/
--qualityThreshold 5
--runThreadN 8
--verbosityDebug 1
```

Individual runs, e.g. to repeat a single experiment were run like this:

```
runVARUS.pl --aligner=HISAT --runThreadN=4 --readFromTable=0 --createindex=1 --verbosity=5 \
  --latinGenus=Prunus --latinSpecies=persica --speciesGenome=Prunus_persica.fa \
  --logfile=Prunus_persica.log 2> Prunus_persica.err
```

##### 1.3 Downloading Manually Selected RNA-Seq Libraries

Manually selected RNA-Seq libraries were downloaded using SRA-Toolkit [Leinonen et al., 2010] as follows ($i is the library ID):

```
fastq-dump $i
```

The argument `--split-files` was appended for paired libraries.

##### 1.4 Aligning Manually Selected RNA-Seq Libraries

All RNA-Seq libraries were aligned using HISAT2 [Kim et al., 2015] version 2.1.0. A genome index was built with `hisat2-build`:

```
hisat2-build genome.fa genome
```

Unpaired RNA-Seq libraries were aligned as follows (${i} is the library ID):

```
hisat2 -p 15 -x genome -U ${i}.fastq -S ${i}.sam 1> ${i}.hisat2.out
```

Paired RNA-Seq libraries were aligned with:

```
hisat2 -p 15 -x genome -1 ${i}_1.fastq -2 ${i}_2.fastq -S ${i}.sam 1> ${i}.hisat2.out
```

SAM format files were subsequently converted to BAM format and sorted by target sequence name using Samtools [Li et al., 2009] version 1.8-20-g4ff8062:

```
samtools view -@18 -bSh ${i}.sam -o ${i}.bam
samtools sort -@18 -n ${i}.bam -o ${i}.s.bam
```

Introns were extracted from BAM files using the AUGUSTUS auxprogs tool `bam2hints` [Hoff and Stanke, 2019]:

```
bam2hints --intronsonly --in=${i}.s.bam --out=${i}.introns
```

Introns from different libraries for the same species were concatenated into a single file per species.

##### 1.5 Running BRAKER

BRAKER [Hoff et al., 2015] version 2.1.3 was executed in order to annotate genomes in a fully automated fashion on the basis of VARUS RNA-Seq alignments and alignments of manually selected RNA-Seq libraries with GeneMark-ES/ET [Lomsadze et al., 2014] version 4.38 and AUGUSTUS [Stanke et al., 2008] version 3.3.2 (intron.hints are either intron hints from manually selected RNA-Seq libaries or sampled by VARUS):

```
braker.pl --genome=genome.fa --hints=hints.introns --softmasking \
  --cores=8 --eval=annot_by_GenomeTools.fixed.gtf
```

The command line option `--fungus` was appended for *Saccharomyces cerevisiae* and *Verticillium dahliae.*

#### 2 Supplementary Figures

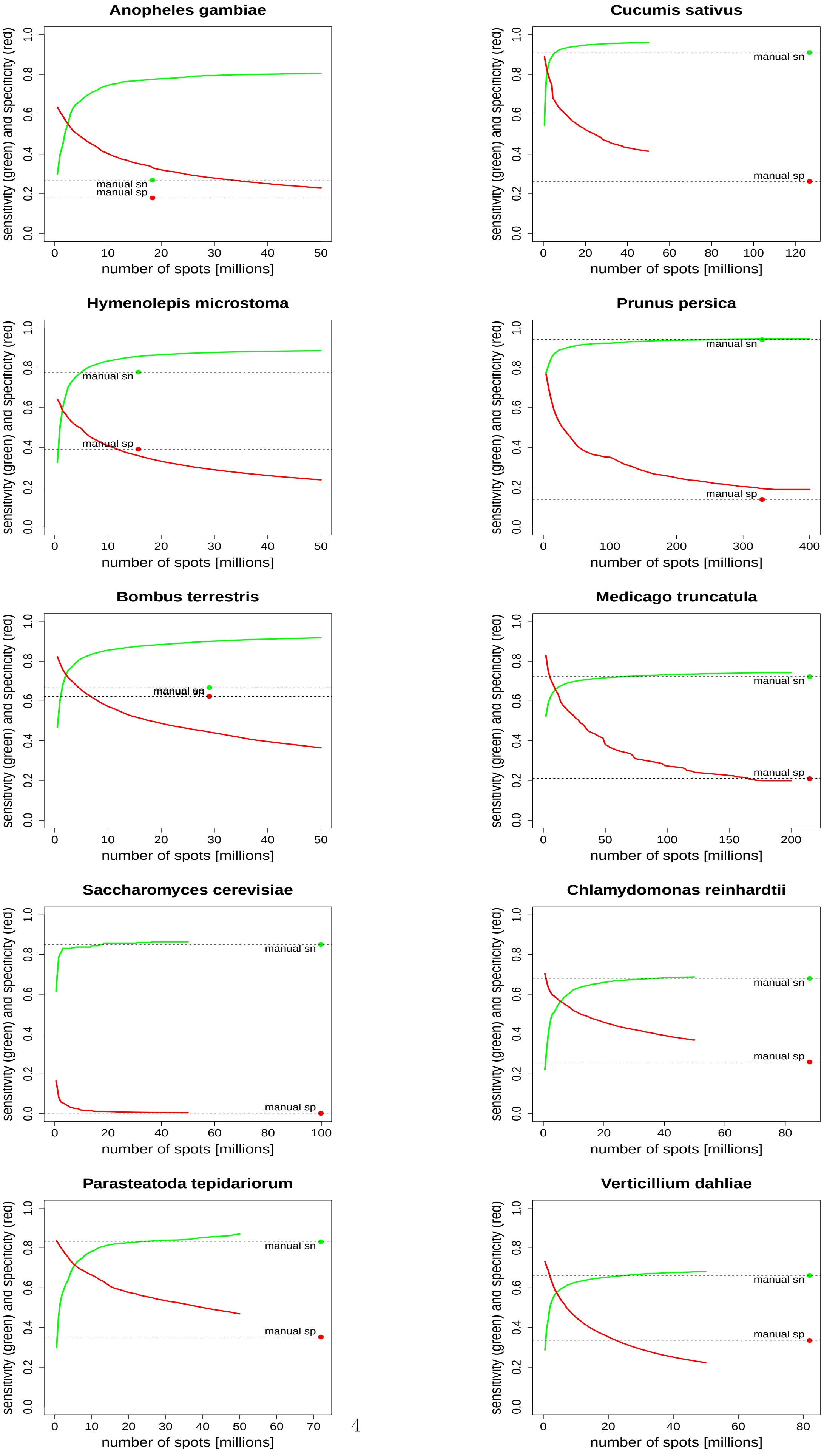

**Supplementary Figure 1:** Intron Accuracy. See Figure 3 of main text for a description.

### 3 Supplementary Tables

| Species | Assembly version | Genome size (bp) |
|---|---|---|
| *Anopheles gambiae* | AgamP3 | 265027044 |
| *Bombus terrestris* | Bter_1.0 | 248654244 |
| *Cannabis sativa* | ASM186575v1 | 585823666 |
| *Chlamydomonas reinhardtii* | v3.0 | 120404952 |
| *Cucumis sativus* | ASM407v2 | 195669205 |
| *Drosophila melanogaster* | r6.07 | 133880608 |
| *Fragaria vesca* | FraVesHawaii_1.0 | 214373013 |
| *Hymenolepis microstoma* | HMIC002 | 182136974 |
| *Medicago truncatula* | MedtrA17_4.0 | 412924424 |
| *Parasteatoda tepidariorum* | Ptep_2.0 | 1445396121 |
| *Prunus persica* | NCBIv2 | 227569171 |
| *Saccharomyces cerevisiae* | R64 | 12071326 |
| *Verticillium dahliae* | ASM15067v2 | 33900324 |

**Supplementary Table 1:** Species names, assembly versions and genome sizes of species used for testing VARUS with BRAKER.

| Species | Annotation source |
|---|---|
| *Anopheles gambiae* | ftp://ftp.ncbi.nlm.nih.gov/genomes/all/GCF/000/005/575/GCF_000005575.2_AgamP3/GCF_000005575.2_AgamP3_genomic.gff.gz |
| *Bombus terrestris* | ftp://ftp.ncbi.nlm.nih.gov/genomes/all/GCF/000/214/255/GCF_000214255.1_Bter_1.0/GCF_000214255.1_Bter_1.0_genomic.gff.gz |
| *Cannabis sativa* | – |
| *Chlamydomonas reinhardtii* | ftp://ftp.ncbi.nlm.nih.gov/genomes/all/GCF/000/002/595/GCF_000002595.1_v3.0/GCF_000002595.1_v3.0_genomic.gff.gz |
| *Cucumis sativus* | ftp://ftp.ncbi.nlm.nih.gov/genomes/all/GCF/000/004/075/GCF_000004075.2_ASM407v2/GCF_000004075.2_ASM407v2_genomic.gff.gz |
| *Drosophila melanogaster* | ftp://ftp.flybase.net/genomes/Drosophila_melanogaster/dmel_r6.07_FB2015_04/gff/dmel-all-r6.07.gff.gz |
| *Fragaria vesca* | ftp://ftp.ncbi.nlm.nih.gov/genomes/all/GCF/000/184/155/GCF_000184155.1_FraVesHawaii_1.0/GCF_000184155.1_FraVesHawaii_1.0_genomic.gff.gz |
| *Hymenolepis microstoma* | ftp://ftp.ncbi.nlm.nih.gov/genomes/all/GCA/000/469/805/GCA_000469805.2_HMIC002/GCA_000469805.2_HMIC002_genomic.gff.gz |
| *Medicago truncatula* | ftp://ftp.ncbi.nlm.nih.gov/genomes/all/GCF/000/219/495/GCF_000219495.3_MedtrA17_4.0/GCF_000219495.3_MedtrA17_4.0_genomic.gff.gz |
| *Parasteatoda tepidariorum* | ftp://ftp.ncbi.nlm.nih.gov/genomes/all/GCF/000/365/465/GCF_000365465.2_Ptep_2.0/GCF_000365465.2_Ptep_2.0_genomic.gff.gz |
| *Prunus persica* | ftp://ftp.ncbi.nlm.nih.gov/genomes/all/GCF/000/346/465/GCF_000346465.2_Prunus_persica_NCBIv2/GCF_000346465.2_Prunus_persica_NCBIv2_genomic.gff.gz |
| *Saccharomyces cerevisiae* | ftp://ftp.ncbi.nlm.nih.gov/genomes/all/GCF/000/146/045/GCF_000146045.2_R64/GCF_000146045.2_R64_genomic.gff.gz |
| *Verticillium dahliae* | ftp://ftp.ncbi.nlm.nih.gov/genomes/all/GCF/000/150/675/GCF_000150675.1_ASM15067v2/GCF_000150675.1_ASM15067v2_genomic.gff.gz |

**Supplementary Table 2:** Sources of reference annotations that were used for assessing gene prediction accuracy of BRAKER1.

| Species | RNA-Seq libraries |
|---|---|
| *Anopheles gambiae* | ERR2192559, SRR5145619, SRR8249138, SRR8249143, SRR8249144 |
| *Bombus terrestris* | ERR047506, SRR098297, SRR5614828 |
| *Chlamydomonas reinhardtii* | DRR059691, SRR057479, SRR094743, SRR6505183, SRR6870054 |
| *Cucumis sativus* | SRR1197971, SRR3872505, SRR517595, SRR7609888, SRR3872503, SRR3872508, SRR6895195 |
| *Drosophila melanogaster* | SRR023505, SRR023546, SRR023608, SRR026433, SRR027108 |
| *Fragaria vesca* | SRR5217589, SRR5217590, SRR5217591, SRR5217592, SRR6320486, SRR6320488, SRR6320491 |
| *Hymenolepis microstoma* | ERR337976 |
| *Medicago truncatula* | ERR1830535, SRR3726837, SRR3938252, SRR7175037, SRR7473103, SRR7772259 |
| *Parasteatoda tepidariorum* | DRR046998, DRR047007, DRR047017, DRR054577, SRR6941355, SRR6941356 |
| *Prunus persica* | SRR5274660, SRR6001701, SRR6001786, SRR6374787, SRR7262551, SRR7469172 |
| *Saccharomyces cerevisiae* | SRR7448271, SRR7696788, SRR7696806, SRR7774370, SRR8261652 |
| *Verticillium dahliae* | SRR2087157, SRR2087158, SRR6130340 |

**Supplementary Table 3:** Manually selected RNA-Seq libraries.

| Species | Exon Sens. | | Exon Spec. | |
|---|---|---|---|---|
| | VARUS | Manual | VARUS | Manual |
| *Anopheles gambiae* | **64.7** | 55.2 | 54.5 | **56.8** |
| *Bombus terrestris* | **78.2** | 71.7 | **70.9** | 67.6 |
| *Chlamydomonas reinhardtii* | **62.1** | 61.6 | 50.0 | **50.1** |
| *Cucumis sativus* | **86.7** | 86.0 | 65.3 | **68.7** |
| *Drosophila melanogaster* | **78.6** | 75.9 | 80.2 | 80.2 |
| *Fragaria vesca* | **84.9** | 83.6 | 62.0 | **66.1** |
| *Hymenolepis microstoma* | **80.3** | 77.9 | 59.5 | **62.8** |
| *Medicago truncatula* | **67.4** | 67.1 | 62.0 | 62.0 |
| *Parasteatoda tepidariorum* | **81.9** | 80.4 | **63.6** | 62.8 |
| *Prunus persica* | 85.1 | 85.1 | 62.2 | **62.9** |
| *Saccharomyces cerevisiae* | **72.5** | 71.2 | **87.4** | 84.8 |
| *Verticillium dahliae* | **64.2** | 63.2 | **64.8** | 64.6 |

**Supplementary Table 4:** Exon level gene prediction accuracy of AUGUSTUS (with hints) trained on the basis of GeneMark-ET predictions within BRAKER1 measured with respect to existing reference annotations. Please note that not all reference annotations are of high quality.
